# A synthetic switch based on orange carotenoid protein to control blue light responses in chloroplasts

**DOI:** 10.1101/2021.01.27.428448

**Authors:** Luca Piccinini, Stefano Cazzaniga, Sergio Iacopino, Matteo Ballottari, Beatrice Giuntoli, Francesco Licausi

**Affiliations:** Plantlab, Institute of Life Sciences, Scuola Superiore Sant’Anna, Piazza Martiri della Libertà 33, 56127, Pis, Italy; Department of Biotechnology, University of Verona, Cà Vignal 1, Strada Le Grazie 15, 37134 Verona; Department of Biology, University of Pisa, Via Luca Ghini 13, 56126, Pisa, Italy; Department of Plant Sciences, University of Oxford, South Parks Road, OX1 3RB, Oxford, UK

## Abstract

**ABSTRACT:** Synthetic biology approaches to engineer light‐responsive system are widely used, but their applications in plants are still limited, due to the interference with endogenous photoreceptors. Cyanobacteria, such as *Synechocystis spp*., possess a soluble carotenoid associated protein named Orange Carotenoid binding Protein (OCP) that, when activated by blue‐green light, undergoes reversible conformational changes that enable photoprotection of the phycobilisomes. Exploiting this system, we developed a new chloroplast‐localized synthetic photoswitch based on a photoreceptor‐associated protein‐fragment complementation assay (PCA). Since *Arabidopsis thaliana* does not possess the prosthetic group needed for the assembly of the OCP2 protein, we implemented the carotenoid biosynthetic pathway with a bacterial β‐carotene ketolase enzyme (crtW), to generate keto‐carotenoids producing plants. The novel photoswitch was tested and characterized in Arabidopsis protoplasts with experiments aimed to uncover its regulation by light intensity, wavelength, and its conversion dynamics. We believe that this pioneer study establishes the basis for future implementation of plastid optogenetics to regulate organelle responses, such as gene transcription or enzymatic activity, upon exposure to specific light spectra.

**One-sentence summary:** Inspired by the light-driven conformational transitions of orange carotenoid proteins of cyanobacteria, we generated a molecular device able to switch its dimeric state in response to blue light.

## INTRODUCTION

Synthetic biology research applies engineering principles to a multidisciplinary approach to decompose natural systems into their essential components and reassemble them to produce novel functions (Gutmann, 2011). Typically, this is aimed at attaining specific applications but can also serve the purpose of understanding naturally occurring biological devices. The synthetic biology framework often taps from (bio)chemistry, genetics and physics to design and assemble biological constructs in an iterative test and optimize process. Due to their modular nature, genes, proteins and metabolites offer a smorgasbord of opportunities, which is mainly limited by the knowledge or predictability of their structure and function, and the possibility to test them. Among the numerous strands of synthetic biology, a particularly prolific area of research concerns the engineering of photoreactive proteins with signaling potential, such as photoreceptors (Olson and Tabor, 2014). Indeed, light quality and quantity, and their dynamics, provide valuable information for cells to integrate with other exogenous and endogenous cues and respond accordingly. Several chimeric constructs have been developed so far, with a focus on the control of neuronal activity by light, thus establishing the field of optogenetics (‘Method of the Year 2010’, 2011).

Light is essential for photosynthetic organisms and thus impacts most of their physiological processes, including development, growth polarity or movement, the internal clock(s) and interaction with the biotic and abiotic environment. Naturally occurring photoreceptors typically consist of a prosthetic chromophore and an apoprotein, responsible for light perception and signal transduction respectively. This generic description applies to most of the photoreceptors found in plants, including tetrapyrrole‐binding phytochromes, flavin‐based cryptochromes and receptors that contain Light–Oxygen–Voltage (LOV) motifs and retinal‐associated rhodopsins (Sineshchekov, Jung and Spudich, 2002; Paik and Huq, 2019). An exception is instead represented by UVR8 and homologs, whose UV light sensing ability is intrinsic to the protein, by virtue of two to four tryptophan residues (Tilbrook *et al.*, 2013). Each photoreceptor perceives to a specific range of visible light spectrum, with some overlaps. Their sensitivity concentrates over red and blue wavelengths, which are especially relevant for photosynthesis, and high‐energy, thus potentially harmful, light. In response to these stimuli, higher plant photoreceptors control flavonoid and chlorophyll biosynthesis, clock entrainment, phototropism, reproductive development (flowering and tuberization), stomatal opening and chloroplast movement (Möglich *et al.*, 2010).

To the repertoire of naturally‐occurring photoreceptors, synthetic biologists have added new ones, exploiting the possibility of domain shuffling across organisms. This has expanded the possibility to fine‐tune molecular processes in response to specific light stimuli. Several plant photoreceptors and their protein partners have been deployed to harness light‐elicited processes in animal cells, for example through engineering Arabidopsis FKF1 (Yazawa *et al.*, 2009), cryptochromes, UVR8 (Kennedy *et al.*, 2010) and phytochromes (Müller *et al.*, 2013). On the other hand, plant optogenetics has lagged behind, due to the extant overlap with a plethora of endogenous photoreceptors. Pioneering efforts in this direction include protein mutagenesis aimed at producing slow‐photocycling phototropin variants (Hart *et al.*, 2019), or engineering green‐light perception through synthetic transcriptional regulators inspired to prokaryotic systems (Chatelle *et al.*, 2018). Main approaches to the engineering of synthetic genetic switches in plant have been recently reviewed (Andres, Blomeier and Zurbriggen, 2019).

Few studies so far sought to engineer new layers of regulation in plant organelles, such as mitochondria and chloroplasts, through synthetic biology approaches, to improve the efficiency of cellular energy factories (Nielsen *et al.*, 2013; Xiang *et al.*, 2020), or to introduce novel methods of transcriptional regulation (Verhounig, Karcher and Bock, 2010). Here we propose the implementation of a new chloroplast‐localized synthetic photoswitch, based on a photoreceptor‐associated protein‐fragment complementation assay (PCA). PCAs exploit the affinity between two peptides fused to split‐protein fragments that, when brought in close proximity, reconstitute the activity of the original protein (Merezhko *et al.*, 2020; Wang *et al.*, 2020).

Our photoswitch system is based on cyanobacterial Orange Carotenoid binding Proteins (OCPs) (Kay Holt and Krogmann, 1981; Kerfeld *et al.*, 2003), which, when activated by blue‐green light, undergo reversible conformational changes that enable phycobilisome photoprotection (Wilson *et al.*, 2006, 2008; Kirilovsky and Kerfeld, 2013). OCPs are water‐soluble proteins, whose structure is mainly composed of a C‐terminal domain (CTD) and an N‐terminal domain (NTD) connected by a flexible linker (Wu and Krogmann, 1997; Kerfeld *et al.*, 2003). The OCP apoprotein non‐covalently incorporates a single keto‐carotenoid molecule, such as canthaxanthin, 3’‐hydroxyl‐echinenone or echinenone, as prosthetic group buried inside the two domains (Punginelli *et al.*, 2009). OCP photoactivation is accompanied by extensive reconfiguration of carotenoid‐protein interactions, with the consequent translocation of the pigment within the protein (Gupta *et al.*, 2015; Leverenz *et al.*, 2015; Bondanza *et al.*, 2020). Photoconversion is possible only when a keto‐carotenoid is buried inside the protein, whilst association with other carotenoids inhibits its photoconversion ability (Punginelli *et al.*, 2009; Wilson *et al.*, 2011).

Here, we report the genetic engineering of *Arabidopsis thaliana* plants to express the chimeric photoswitch based on OCPs and the NanoLuc luciferase protein, together with a bacterial enzyme for keto‐carotenoid biosynthesis to generate the necessary prosthetic group (Ruiz-Sola and Rodríguez‐ Concepción, 2012; Nisar *et al.*, 2015; Bai *et al.*, 2017). We also describe the application of transient transformation systems to fully characterise the dynamics of such synthetic construct to establish the bases of its future applications.

## RESULTS

### Design of a photo‐switchable device based on OCP properties

To engineer a light‐dependent molecular switch to be orthogonally active in plant plastids, we focused on the class of the cyanobacterial Orange Carotenoid‐binding Proteins (OCPs). Among the three known paralogous families of OCPs (OCP, OCP2 and OCPX), we reckoned the OCP2 family to fit better the rational design of a synthetic photoreceptor. Indeed, its faster conversion kinetics, its monomeric state, and its ability to revert to the dark‐adapted state (OCP2°) in the absence of a helper protein (Bao *et al.*, 2017) reduce the number of components required for the photoswitch to function. Thus, we selected the OCP2 coding sequence (CDS) from the cyanobacterium *Fischerella thermalis* and separated its N‐terminal domain (NTD, 165 aa) and C‐terminal domain (CTD, 131 aa). *F. thermalis* was chosen as the source organism as one of the few known to encode for an OCP2 protein as the sole OCP sequence in its genome (Bao *et al.*, 2017). Moreover, being *F. thermalis* a themophilic organisms (Alcorta *et al.*, 2019), proteins encoded by its genome likely have higher stability compared to the case of non‐thermophilic ones. To test whether these modules reconstitute a complex capable of light‐dependent regulation in angiosperm cells, we generated a synthetic construct that couples a luminescent output with dimerisation (Fig. 1). In our design, we exploited the affinity and light‐driven separation of the two OCP2 modules for a protein complementation assay, based on a NanoLuc complementation assay developed in mammalian cells and called NanoBit (Dixon et al. 2016).

**Figure 1.**
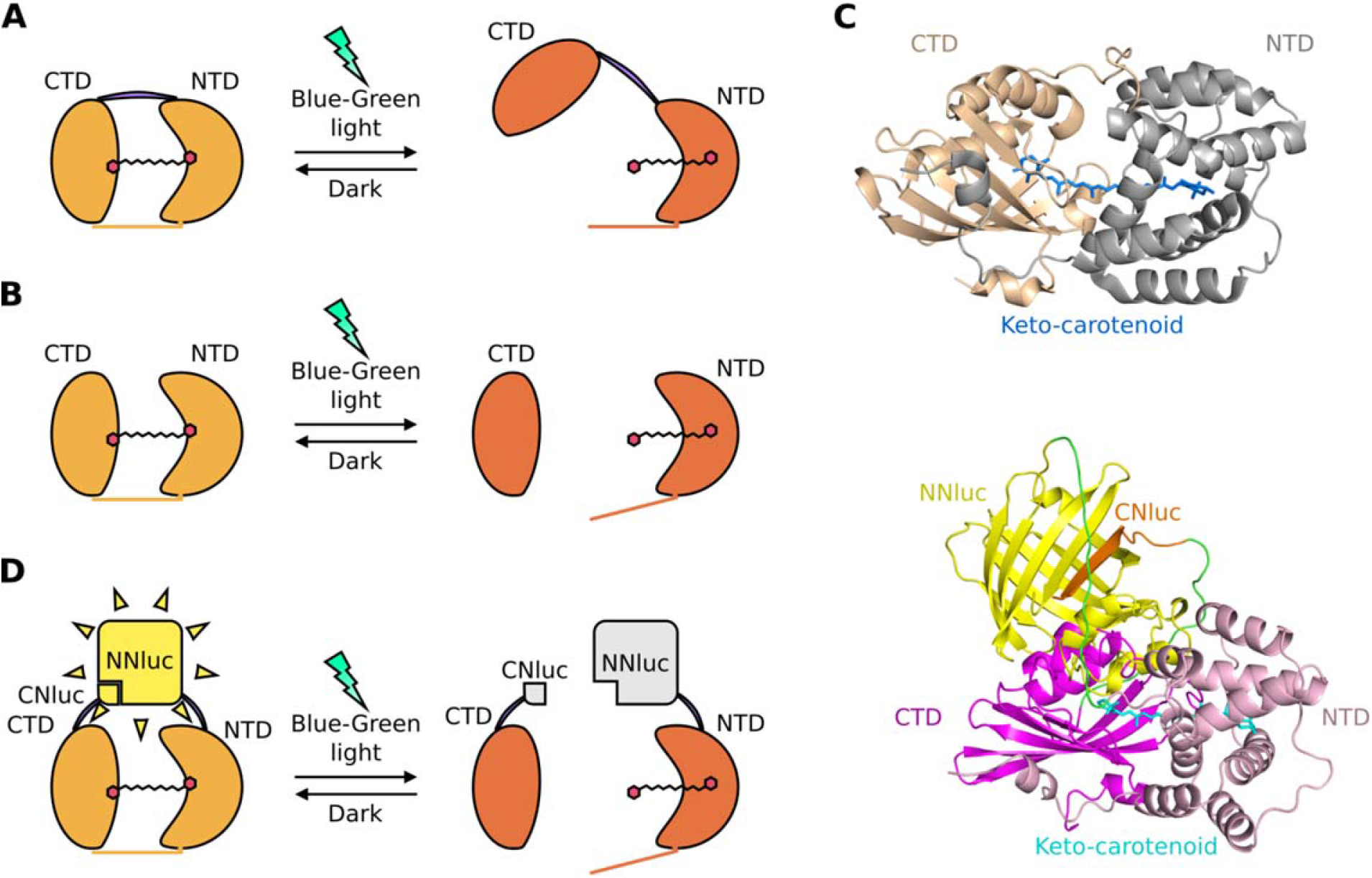
Rational design of a blue‐green photoswitch based on OCP2 modules and the NanoLuc complementation system (split‐NlucOCP2). A, Scheme of the light‐induced native OCP2 protein dynamics: dark state (left), called OCP2 orange (OCP2°), and light‐induced state (right), known as OCP2 red (OCP2^R^). B, N‐Terminal Domain (NTD) and C‐Terminal Domain (CTD) interactions, when expressed as separate peptides, in darkness (left) or under stimulating light. C, *In silico* prediction of dark OCP2 (top) and dark split‐NlucOCP2 (bottom) 3D structures. D, Scheme of the split‐NlucOCP2 photoswitch. In the dark, the two OCP2 modules interact with each other, thus bringing the two NanoLuc fragments in close proximity and complementing NanoLuc; blue‐green light leads to a separation of the two modules, suppressing NanoLuc activity.

We fused each domain with one of the two fragments of the split‐NanoLuc protein: the N‐terminal (NNluc, 159 aa) and the C‐terminal (CNluc, 11 aa) fragments. In this way, we generated two chimeric constructs which, together, constitute a split‐NlucOCP2 photoswitch. In the wild‐type OCP2 protein, the two domains associate by means of a flexible linker and by the non‐covalent bonds that a single keto‐ carotenoid molecule establishes after its incorporation inside the OCP2 protein (Wu and Krogmann, 1997; Kerfeld *et al.*, 2003) (Fig. 1A‐B). Instead, in our synthetic construct, the OCP2 modules are kept together exclusively by their interactions with the keto‐carotenoid (Fig. 1B). Since the strength of these interactions depends on the light‐dependent translocation of the keto‐carotenoid inside the OCP2 structure (Gupta *et al.*, 2015; Leverenz *et al.*, 2015; Bondanza *et al.*, 2020), we expected the NTD and the CTD to associate in the dark, whereas exposure to blue‐green light would bring them apart.

Specifically, we fused the NNluc to the C‐terminus of the NTD module (NTD‐NNluc), and the CNluc to the N‐terminus of the CTD module (CNluc‐CTD). This conformation was selected following an *in silico* prediction based on the 3D structures of NanoLuc (Protein Data Bank (PDB) ID 5IBO) and the OCP from *Limnospira maxima* (PBD ID 5UI2), which shares the strongest sequence similarity with FtOCP2 among the OCPs whose structure has been experimentally determined. This model suggested these fusions as the best option to bring the NanoLuc modules in close proximity in the closed (OCP°) conformation (Fig. 1C).

We predicted that the interaction between the NTD and the CTD in the dark would reconstitute the enzymatic NanoLuc activity and therefore produce a luminescent output upon addition of the furimazine substrate to plant cell extracts (Hall *et al.*, 2012; Dixon *et al.*, 2016). On the other hand, we expected blue‐green light exposure to promote detachment of the CTD from the NTD, resulting in the disruption of the NanoLuc complementation (Fig. 1D). The low affinity between the NNluc and the CNluc themselves (Dixon et al. 2016), allowed us to rule out that the reconstitution of the NanoLuc activity would only be due to their interaction.

### Generation and characterization of canthaxanthin‐producing Arabidopsis transgenic plants

Both the interaction and the photoconversion of the OCP2 modules depend on the presence of specific keto‐carotenoids (canthaxanthin, 3’‐hydroxyl‐echinenone or echinenone) (Punginelli *et al.*, 2009). For our synthetic photoswitch to be functional, one of the two modules, usually the NTD (Lechno-Yossef *et al.*, 2017), should be conjugated with a single molecule of the proper keto‐carotenoid. Therefore, we generated stable transgenic Arabidopsis lines expressing the β‐carotene ketolase (crtW) enzyme from *Agrobacterium aurantiacum*, which catalyzes canthaxanthin biosynthesis from β‐carotene (Fig. 2A) via the addition of a carbonyl group to carbon 4 and 4’ of the substrate (Misawa *et al.*, 1995; Choi *et al.*, 2005; Bai *et al.*, 2017). It is important to note that the crtW enzyme is also involved in the biosynthetic pathway of another ketocarotenoid, astaxanthin, by using zeaxanthin as substrate. Moreover, canthaxanthin itself can be hydroxylated by an endogenous β‐hydroxylase enzyme, leading to astaxanthin production (Zhong *et al.*, 2011). Since carotenoid biosynthesis in higher plants occurs in plastids (Nisar *et al.*, 2015), we fused the sequence of the crtW enzyme with a previously tested plastid localization sequence (PLS) from ribulose 1,5 bisphosphate carboxylase of *Pisum sativum* (Fig. 2B), and placed this *PLS‐crtW* transgene under the control of the constitutive *Cauliflower mosaic virus* (CaMV) 35S promoter.

**Figure 2.**
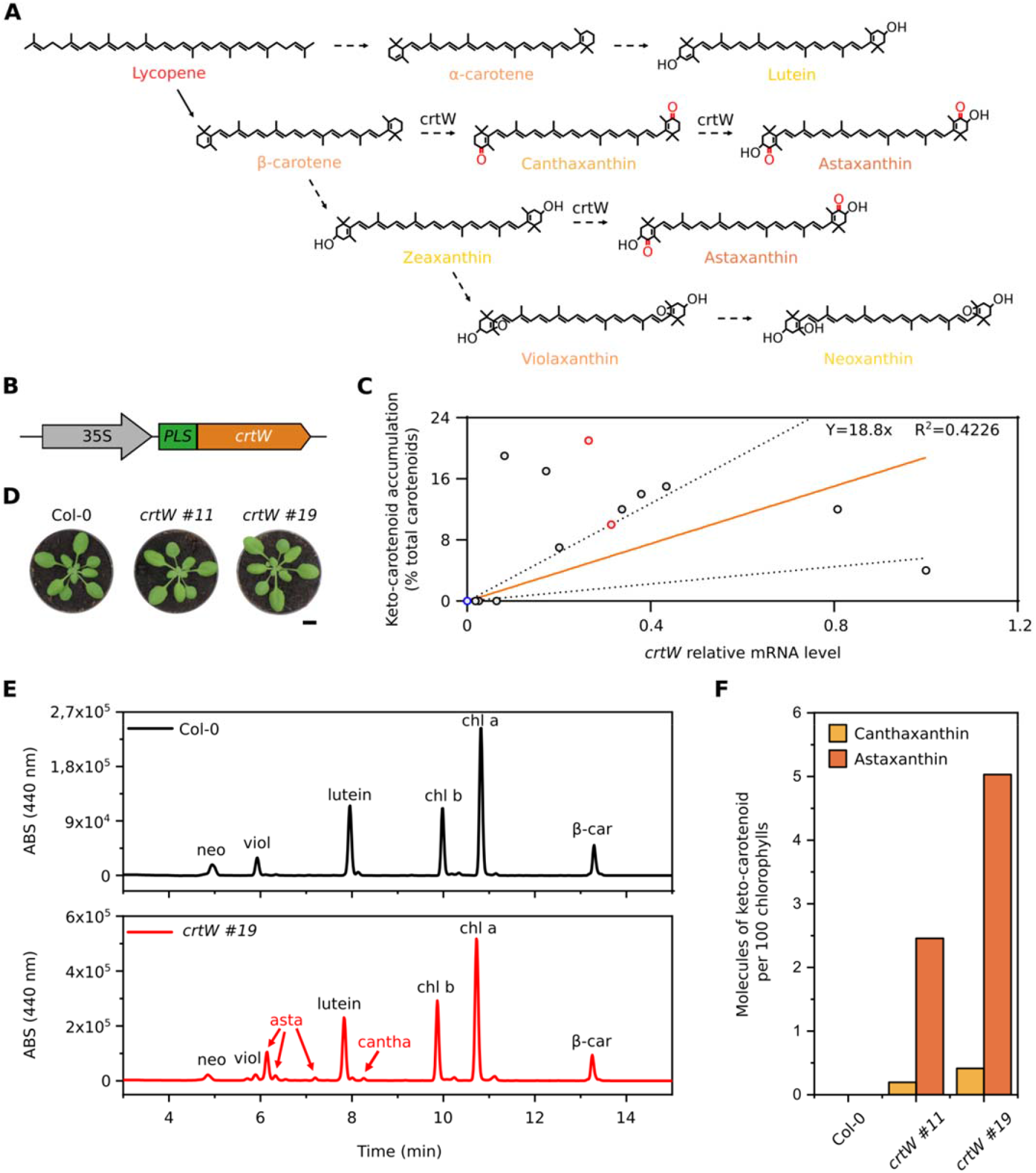
Pigment content analysis of keto‐carotenoid *Arabidopsis thaliana* transgenic lines. A, Overview of the biosynthetic carotenoid pathway after the addition of the crtW enzyme; dotted arrows represent multiple enzymatic steps. B, Scheme of the synthetic construct for canthaxanthin production: *Cauliflower mosaic virus* 35s promoter (CaMV35s); Plastid localization sequence (PLS); β‐carotene ketolase (crtW). C, Relative *crtW* mRNA levels and their corresponding keto‐carotenoid content in 13 independent transgenic lines (black and red circles) and one Col‐0 (blue circle); red circles represent the two lines (#11 and #19) chosen for the following experiments. mRNA levels in the graph are relative to the transgenic line with the highest expression, set as 1. The orange line represents the curve fitted to the data with linear regression, and the dotted lines are the 95% prediction confidence intervals. The R squared (R^2^) and the equation of the curve are shown. D, Phenotypic comparison between Arabidopsis Col‐0 plants and two independent *crtW* transgenic lines. E, HPLC analysis for canthaxanthin and astaxanthin detection in a *crtW* line (red line), compared with a Col‐0 plant (black line); neoxanthin (neo), violaxanthin (viol), astaxanthin (asta), canthaxanthin (cantha), chlorophyll b (chl b), chlorophyll a (chl a), β‐carotene (β‐car). F, Quantitative measurement of canthaxanthin and astaxanthin, represented as molecules of keto‐carotenoid per 100 molecules of chlorophylls.

Following positive selection on kanamycin, 21 putative transformants were isolated. To evaluate the actual expression of the transgene, we measured the *PLS‐crtW* transcript level through real time quantitative PCR (RT‐qPCR) and correlated it to the content of ketocarotenoids, as measured by High Performance Liquid Chromatography (HPLC) (Fig. 2C). All selected *35S:PLS‐crtW* plants actively expressed the transgene with extensive variability. Out of 13 lines analysed, 10 expressed detectable levels of keto‐carotenoids, ranging from 4% to 21% of the total pigment content (Fig. 2C). Astaxanthin was the most abundant keto carotenoid accumulated, but all 10 lines also contained the OCP‐binding canthaxanthin. Keto carotenoids were synthesized at the expense of the β‐β xanthophyll neoxanthin and violaxanthin, which were reduced with respect to the parental lines. Zeaxanthin was not detected in any plant in the light regime used in this analysis. Total carotenoid content was not different between the genotypes.

Two independent lines with high and intermediate keto‐carotenoids content (Fig. 2C, red circles, and Fig. 2D) were further analyzed (Fig. 2E and Fig. 2F). At a visual inspection, the two transgenic lines showed a slightly brown pigmentation compared to the wild‐type, which was only partially confirmed by a phenotyping analysis based on RGB imaging of the plants through a commercial phenotyping machine. Comparing Col‐0, *crtW #11* and *crtW #19* genotypes at 17, 21 and 24 days after germination, we found no significant difference in the average size of the plants, and comparison of the three components of the RGB images (Red, Green and Blue) showed a statistically significant increase in the red component between the 17 and 21 days old Col‐0 and *crtW #19* (Supplemental Fig. S1). Considering the gathered evidence collectively, we concluded that the implementation of the endogenous carotenoid biosynthetic pathway allowed the production of keto‐carotenoids in Arabidopsis.

### Testing of OCP2‐based photoswitch in plant protoplasts

We generated plasmids carrying the NTD‐NNluc and CNluc‐CTD transgenes, each equipped with a plastid localization signal (PLS), to be expressed under control of the 35S CaMV promoter, and we tested the functionality of the photoswitch system in transient assays in mesophyll protoplasts. We started by investigating the activity of the constructs in protoplasts isolated from either Col‐0 or *crtW* plants, to assess whether keto‐carotenoids were required for the output of the system, according to our rational design.

A first proof‐of‐principle experiment was performed to evaluate the responsivity of the system to the blue‐green light stimulus and its basal activity. All possible combinations (Fig. 3A) of the two modules were transformed in *crtW* protoplasts. Sixteen hours after the transformation, dark‐incubated protoplasts were treated under blue‐light (465‐480 nm, 350 μmol m^−2^ s^−1^) for 1 hour, while control samples were maintained in the dark. Protoplasts transformed with PLS‐CNluc‐CTD or PLS‐NTD‐NNluc alone showed no luminescent output in the dark, confirming that the two NanoLuc fragments did not retain any basal enzyme activity (Fig. 3A). Those transformed with both modules were instead analysed for light response. High luminescence output in dark‐treated protoplast was suggestive of effective complementation and reconstitution of a dark‐adapted state in the OCP2 complex. The output was instead significantly lower in illuminated protoplasts, indicating the detachment of the OCP2 modules, as expected, with consequent disruption of the enzymatic activity (Fig. 3A). Wild‐type protoplasts had identical output upon dark or light treatment, suggesting that light is unable to trigger the structural changes in the OCP2 complex in the absence of the proper keto‐carotenoids. The residual activity recorded in the wild‐type background pointed at the occurrence of some extent of unspecific interaction between the two photoswitch modules, likely due to the affinity of the OCP2 fragments. Remarkably, however, blue‐green light treatment in the *crtW* background cut the output down to the level associated with unspecific module interaction (Fig. 3A), suggesting that the keto‐carotenoid dependent changes that are typical of the native OCP2 protein were fully recapitulated in our split configuration. Next, we investigated the effect of a different subcellular localization of the photoswitch modules. In plants, carotenoids are mainly concentrated inside plastids, and no specific or generic carotenoid export system is known (Sun *et al.*, 2018). To exclude our photoswitch from plastids, we generated two new versions of the CNluc‐CTD and NTD‐NNluc modules devoid of PLS. The new modules were again tested both in *crtW* and Col‐0 protoplasts, using the plastid localized photoswitch as positive control for the NanoLuc activity. We found that the luminescent signal produced in the cytosol‐localized photoswitch modules was comparable to that of untransformed protoplasts (Fig. 3B), confirming that our photoswitch can only be active in carotenoid‐containing organelles.

**Figure 3.**
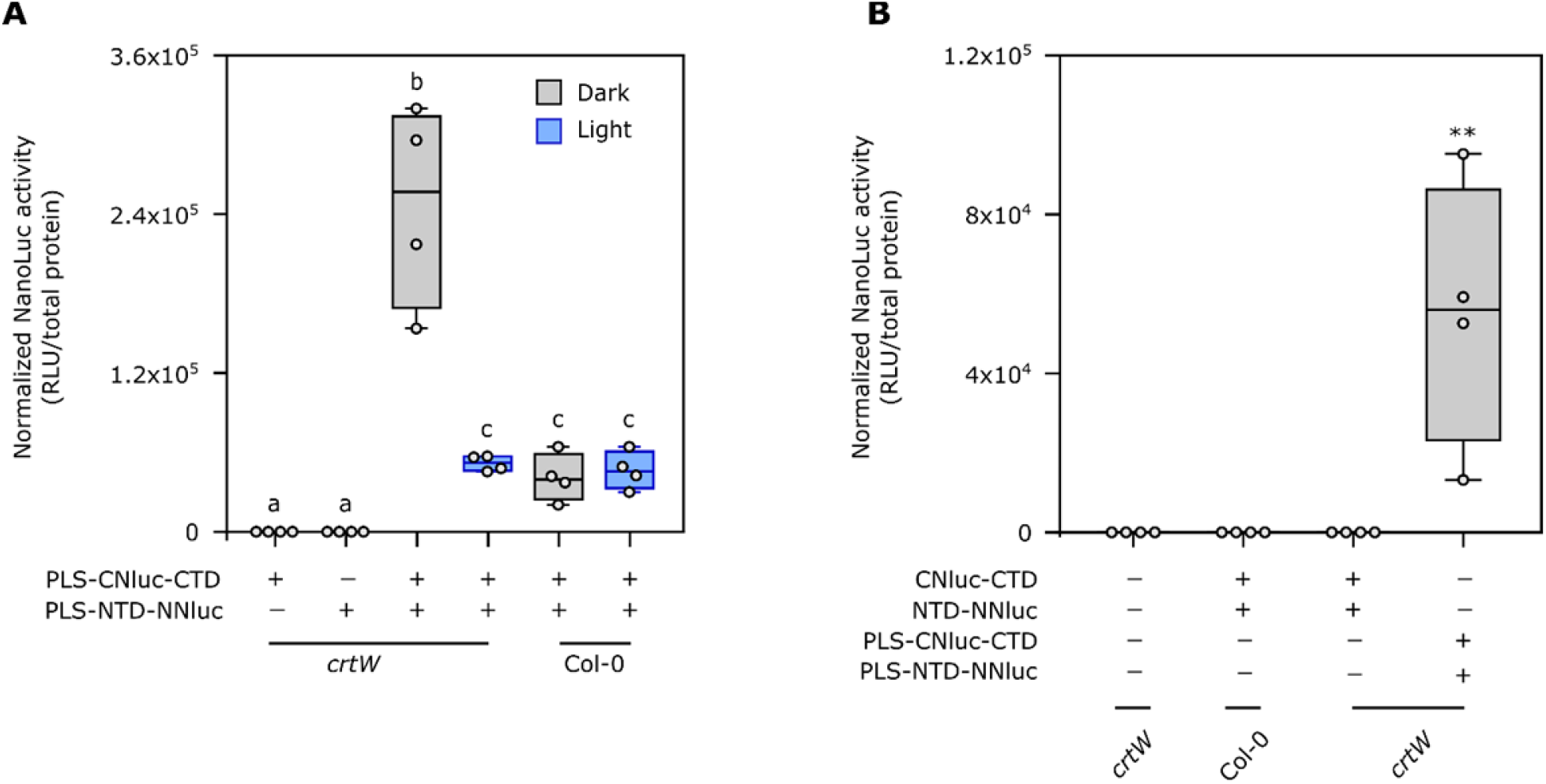
Activity of the synthetic photoswitch in a transient protoplasts assay. A, Light response in Arabidopsis Col‐0 and transgenic *crtW* ‐isolated protoplasts. Different letters indicate statistical differences (P ≤ 0.05) calculated from one‐way ANOVA (n=4) followed by Tukey’s post‐hoc test. “Light” indicates treatment under blue‐green light (465‐480 nm, 350 µmol μm^‐2^ s^‐1^) for 1 h. B, Comparison between plastidial and non‐plastidial localization of photoswitch components. Protoplasts were treated with either dark or blue light (350 µmol μm^‐2^ s^‐1^) for 1 hour. Asterisks indicate statistical differences (P ≤ 0.01, n=4). In the box plots, dots represent single data points, the black line marks the median, and the box indicates the interquartile range (IQR). Whiskers extend to data points below 1.5 X IQR away from the box extremities.

### Characterization of split‐NlucOCP2 light‐responsiveness and reversion

Once we found out the basal requirements of our photoswitch, we performed a full characterization of its dynamics, to describe how its activity could be modulated by the light. First, to understand the kinetic of the light‐induced structural changes in the photoswitch, we treated the transformed *crtW* protoplasts for different periods with white light (350 μmol m^−2^ s^−1^), from 5 to 80 minutes, after dark incubation. Photoconversion of a significantly high amount of the complex occurred as early as 5 mins after light exposure and increased further with time (Fig. 4A). We selected 20 mins light treatment for further analysis, since this data point was associated with a low variability among the biological replicates. Next, we investigated whether light quality is relevant for the regulation of our photoswitch activity, and which wavelength range is the most effective in triggering the switch. Treatments with a tuneable led lamp emitting red (615‐630 nm), green (515‐530 nm), blue (465‐480 nm) or white light were compared with dark treated samples (Fig. 4B). We observed that the output was significantly different between the dark treatment and the whole set of different wavelengths: blue and green light exerted the strongest effect, while red light only halved the luminescent output. White light was not as effective as blue light to switch off the system, suggesting that saturated blue light is required to achieve the maximum photoswitch performance.

**Figure 4.**
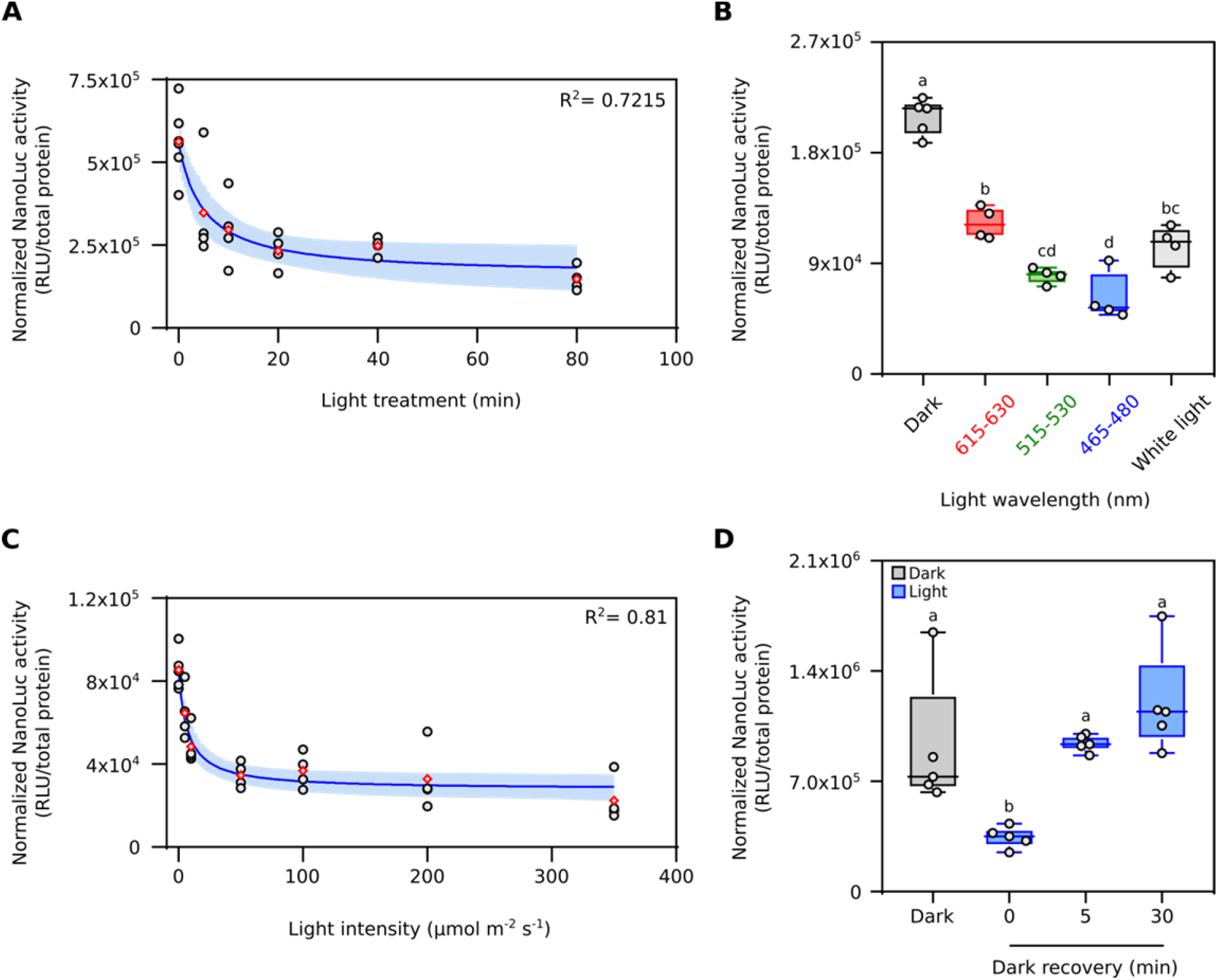
Characterization of the split‐NlucOCP2 photoswitch in *Arabidopsis thaliana* protoplasts. A, Impact of different lengths of light exposure (white light, 350 µmol m^‐2^ s^‐1^) on photoswitch output. Data points correspond to 0, 5, 10, 20, 40 and 80 minutes. B, Effect of different wavelengths on the output of the photoswitch, as compared to darkness. Red, green, blue or white light was supplied for 20 minutes at 350 µmol m^‐2^ s^‐1^). C, Output modulation under increasing light intensity. Protoplasts were exposed to 5, 10, 50, 100, 200, or 350 µmol m^‐2^ s^‐1^ blue light for 20 minutes after 16 h dark incubation (initial data point). D, Test of the reversibility of the photoswitch. After light treatment (20 minutes, 50 μmol m^‐2^ s^‐1^, blue light), the samples were incubated for 5 or 30 min in the dark before sampling. In the box plots, dots represent single data points, the black line marks the median, and the box indicates the interquartile range (IQR). Whiskers extend to data points that are below 1.5 X IQR away from the box extremities. Different letters indicate statistical differences (P ≤ 0.05) calculated from one‐way ANOVA followed by Tukey’s post‐hoc test. In dot plots, data fitting through non‐linear regression analysis is shown with blue continuous curves, generated using the GraphPad Prism built‐in equation for [Inhibitor] vs. response model. Shaded bands indicate the predicted 95% confidence interval. Black dots represent single data points and red rhombuses represent mean values. The R squared (R^2^) of the curves are included in the graphs.

Having established the optimal treatment duration (20 minutes) and light spectrum (blue) to elicit photoswitch output, we moved on to analyse the effect of light intensity on our system, testing the range from 5 to 350 μmol m^−2^ s^−1^ (Fig. 4C). We could measure significant signal decrease already at 10 μmol m^−2^ s^−1^. The response was saturated at 50 μmol m^−2^ s^−1^. Finally, we investigated the possibility that our synthetic photoswitch could revert from the dissociated to the dimeric form when light exposure ceases, as happens for heterodimeric OCP2 complexes *in vitro* (Lechno-Yossef *et al.*, 2017). We shifted protoplast samples, previously exposed to 20 min saturating blue light (50 μmol m^−2^ s^−1^), to darkness for different amounts of time. We found out that the split‐NlucOCP2 had the capacity to revert quickly to the dark‐adapted conformation after the treatment, since we measured full recovery of the signal as early as 5 min into dark incubation (Fig. 4D).

## DISCUSSION

In this work, we successfully exploited the properties of a cyanobacterial Orange Carotenoid Protein to generate a synthetic photoswitch meant to be active in plant chloroplasts (Fig. 3B). We were able to draw a complete characterization of the prototype version of our synthetic photoswitch, split‐NlucOCP2, in living plant cells, thanks to a strategy based on protoplast transformation. Synthetic biology endeavours in plants are complicated by non‐negligible hurdles associated with genetic engineering, such as long timeframes for the generation of transgenic individuals, low rates of homologous recombination and, in some cases, cumbersome procedures for plant genetic transformation. However, populations of isolated cells, obtained from the enzymatic digestion of cell walls from plant tissues (typically young, poorly lignified leaves), can be adopted as an easy and convenient alternative to implement fast, combinatorial and high‐throughput tests for newly designed synthetic devices (Yoo, Cho and Sheen, 2007; Pouvreau *et al.*, 2020). We therefore opted for this strategy to characterize the modules of split‐NlucOCP2.

As according to our design (Fig. 1D), this system was inhibited by light inputs; the stimulus was sufficient to promote the maximum extent of monomerization of the system, indicated by the activity observed in the Col‐0 background (Fig. 3A). Another important feature of the photoswitch was its ability to revert spontaneously to the heterodimeric (dark‐adapted) state after the light‐induced monomerization (Fig. 4D). The extent of output inhibition by light was directly proportional to the treatment duration and intensity (Fig. 4A and C). Altogether, this evidence let us conclude that split OCP2 strategies are viable to design fast‐responding and quickly reversible photoswitches, characterized by a continuous response to light and a high dynamic range. The blue and green ranges of the light spectrum were the most effective ones in switching off the response (Fig. 4B). Since white light contains blue, green and red wavelengths, we were not surprised by its effect on the switch (Fig. 4B); the ability of red light to inactivate our photoswitch, however, was unexpected, since OCP2 has an absorption peak in the blue‐green range at 474 and 502 nm and cannot be photoactivated by red‐light (Bao *et al.*, 2017). We may speculate that this effect is caused by differences in the spectral properties of both the keto‐carotenoid, due to the protein environment, and the protein, since we are expressing a split and engineered version of it.

Two features proved to be essential for the synthetic photoswitch to control luciferase activity: firstly, chloroplasts needed to produce the required keto‐carotenoid prosthetic group and, secondly, the two photoswitch modules required plastidial localization. Thus, we had to implement a canthaxanthin biosynthetic pathway by expressing an exogenous β‐carotene ketolase (crtW) enzyme, which catalyzes canthaxanthin biosynthesis (Fig. 2). Indeed, when protoplasts expressed cytosol‐localized modules (Fig. 3A) or were unable to produce keto‐carotenoids (Fig. 3B), the switch was non‐functional. Dimerization and photoativation of the photosowitch could be related to either canthaxanthin or astaxanthin, which in both cases were recently demonstrated to be bound by OCP2, in the case of heterologous expression of this subunit in the green alga *Chlamydomonas reinhardtii* engineered for producing keto‐carotenoids (Pivato et al resubmitted). In the keto‐carotenoid producing plants, we found that astaxanthin was reproducibly more abundant than canthaxanthin (Fig. 2F and Table 1). This is not entirely unexpected/surprising, given previous demonstrations that the addition of the *crtW* enzyme in plant provides only one pathway towards canthaxanthin, whereas more pathways converge towards astaxanthin production (Bai *et al.*, 2017), one of which using canthaxanthin as a substrate. Comparison of the composition of the carotenoid pool in *ctrW* and wild‐type plants showed that keto‐carotenoids were synthesized at the expense of other β‐β xanthophylls. As shown in Table 1, neoxanthin, β‐carotene and violaxanthin present lower average values in keto‐carotenoids producing lines, even though only the latter shows a statistical significance. This was expected, because β‐carotene and zeaxanthin are substrate of the crtW enzyme, but also precursors of both neoxanthin and violaxanthin (Fig. 2A). On the other hand, ε‐β xanthophyll lutein, whose biosynthesis follows a different branch of the pathway, (Ruiz-Sola and Rodríguez‐Concepción, 2012; Bai *et al.*, 2017) was unaffected (Fig. 2A).

**Table 1.**
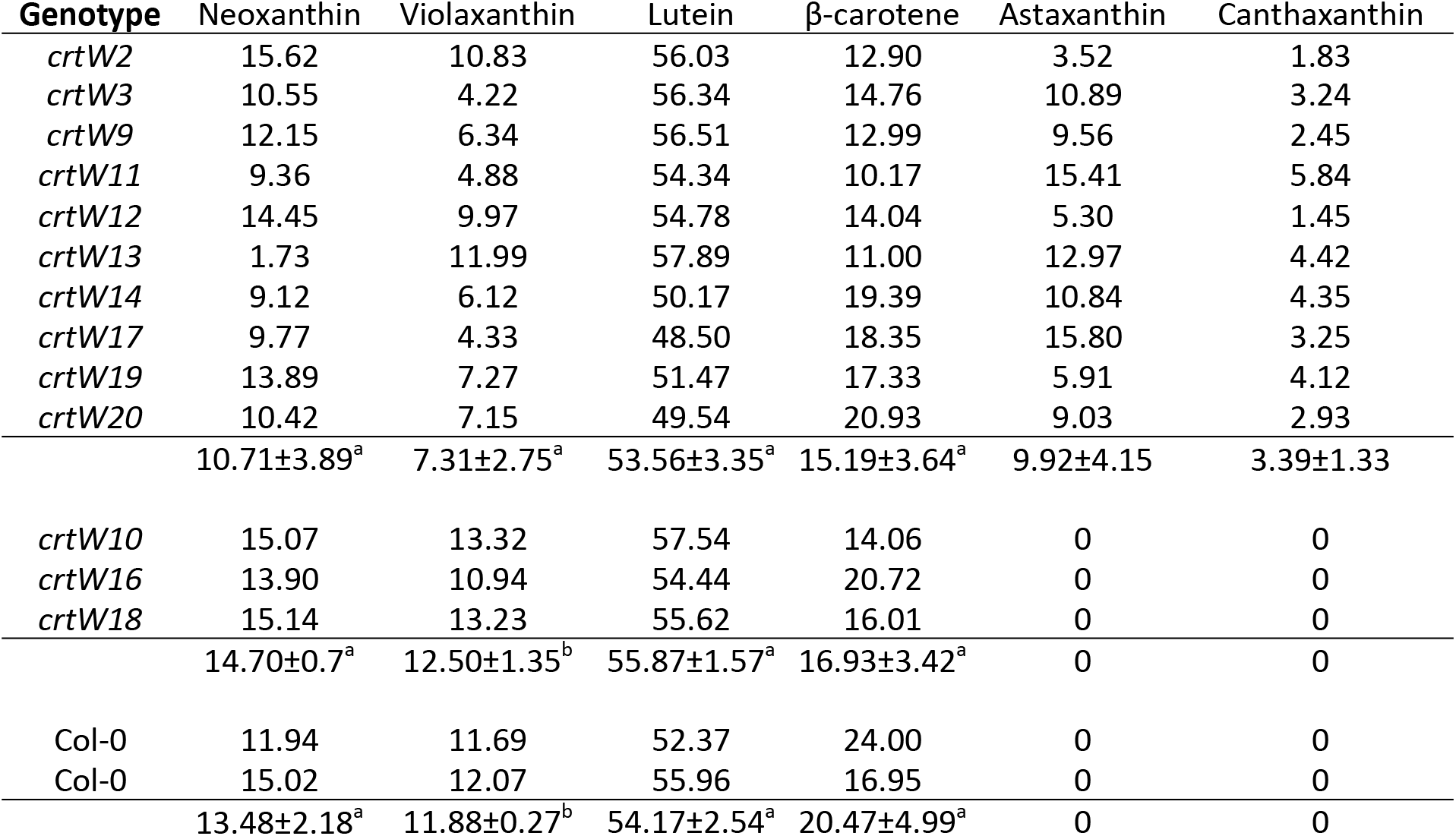
Comparison of the carotenoid content in crtW and wild‐type Arabidopsis plants. Data are percentages of each pigment relative to the total carotenoid content per each line. Average and standard deviation are shown per each group of lines. A Student T‐test was used to assess significant differences between *crtW* and Col‐0 genotypes. Different letters between rows represent differences in carotenoid content (p<0.05).

Light exposure could not abolish OCP2‐nanoLuc signal entirely, even after several hours. The signal still present in light‐treated *crtW* protoplasts (Fig. 3A) can be explained by two non‐mutually exclusive hypotheses: that the treatment could not separate the whole pool of interacting NTD‐CTD modules, or that other carotenoids, such as β‐carotene, may mediate modules interaction, although without the ability to photoconvert (Punginelli *et al.*, 2009; Wilson *et al.*, 2011). Recently, *Fischerella thermalis* OCP2 was shown to bind neoxanthin, loroxanthin and lutein when heterologously expressed in *Chlamydomonas reinhardtii* (Pivato et al. resubmitted). The latter scenario could also explain the amount of signal we measured in dark treated wild‐type protoplasts, which was not affected by the light treatment (Fig. 3A). Moreover, the complete absence of signal observed in the cytosol‐localized photoswitch suggests that none of the carotenoids able to conjugate with OCP2 were present outside the chloroplast in sufficient amounts to associate with the photoswitch (Fig. 3B).

The success of this initial study lays the foundations for different ways in which our system could be exploited in the future. Similar to NanoLuc, the OCP2 modules can be fused to a bipartite enzyme or structural protein whose function is favoured under dark conditions and/or not required or even detrimental when cells are illuminated. Alternatively, the OCP2 modules could serve for reversible association of an inhibitor protein, thus leading to positive regulation upon light exposure and repression in darkness. While potentially presenting a limitation for use in plants that are not able to synthesize keto‐carotenoids, our system could be exploited not only for its photo‐induced activity, but also as a carotenoids biosensor: when carotenoids (β‐carotene, zeaxanthin) are present in the cellular environment, a small percentage of the modules is expected to dimerize in a stable manner; instead, if in case of canthaxanthin, 3’‐hydroxyl‐echinenone or echinenone accumulation, the dimerization rate will be higher and reversible under light treatment (Fig. 3A). Alternatively, this photoswitch strategy might be applied in microalgae, where different species accumulate keto‐carotenoids or can be successfully engineered to express them (Perozeni *et al.*, 2020).

In summary, here, we demonstrated that plant organelles can be equipped with an orthogonal synthetic photoswitch, based on a cyanobacterial light‐responsive protein and its cognate keto‐carotenoids. The behavior of this synthetic construct led us to conclude that, the structural modularity of OCP, together with its light‐driven dissociation (Leverenz *et al.*, 2014; Lechno-Yossef *et al.*, 2017), makes it a suitable tool for optogenetic applications (Dominguez‐Martin and Kerfeld, 2019). Selecting biological modules from unrelated species grants the highest probability that cross‐interactions between components and endogenous pathways are kept to a minimum, and we consider particularly interesting the ability of OCPs to be triggered by the green range of the light spectrum, considering that it should not interfere and stimulate the most common photoreceptors already known inside the plant systems, ensuring the orthogonality of the system. We believe that this pioneer study establishes the basis for future implementation of plastid optogenetics to regulate organelle responses, such as gene transcription or enzymatic activity, upon exposure to specific light spectra.

## MATERIALS AND METHODS

### Design of synthetic DNA sequences and plasmid assembly

Sequences were designed, codon‐optimized for A. *thaliana* expression and synthesized as DNA strings using the GeneArt service (Thermo‐Fisher Scientific). The sequences of the synthetic constructs devised for this work is provided in Supplemental Information S1. Directional cloning of DNA fragments was performed using the pENTR™ Directional TOPO^®^ Cloning Kits (Thermo‐Fisher Scientific). For the OCP2 modules we took the *Fischerella thermalis* OCP2 sequence as reference (accession WP_009459388). The non‐chloroplast localized fragments were amplified with the Phusion™ High‐fidelity DNA polymerase (Thermo‐Fisher Scientific) using the following primer couples: NTD‐NNluc‐Fw (CACCATGTCCTTCACCATCGAG) with NTD‐NNluc‐Rv (TCAGCTGTTGATGGTCACTCTG) and CNluc‐CTD‐Fw (CACCATGGTGACCGGGTACA) with CNluc‐CTD_Rv (TCACTGGATGAATCCCATGTTG). The resulting entry vectors were then recombined into destination vectors via Gateway™ LR Clonase™ II Enzyme mix (Thermo‐Fisher Scientific), to produce the desired expression constructs. A 35S:*PLS‐crtW* construct for constitutive transgene expression in plants was obtained by recombination of the proper entry vector, containing the *crtW* coding sequence from *Agrobacterium aurantiacum* (*Paracoccus sp.* N81106), with the binary vector pK7WG2 (Karimi, Inze and Depicker, 2002). The individual modules PLS‐NTD‐NNluc, PLS‐CNluc‐CTD, NTD‐NNluc and CNluc‐CTD were, instead, recombined with the plasmid p2GW7 (Karimi, Inze and Depicker, 2002), suitable for transient expression in Arabidopsis protoplasts.

### Protein structure modeling

The three‐dimensional structure of CNluc‐CTD and NTD‐NNluc was generated according the following procedure: the amino acid sequences of both CNluc‐CTD and NTD‐NNluc were aligned to the template sequences of Orange Carotenoid Protein of *Limnospira maxima* (PDB: 5UI2) (Kerfeld et al. 2003) and the synthetic luciferase NanoLuc (PDB: 5IBO) (Lovell et al., unpublished) using MultAlin (Corpet, 1988) and the resulting alignment used as input file for modeling using Modeller(Webb and Sali, 2016). Figure rendering was performed with the PyMol software (DeLano, 2020).

### Plant materials and growth conditions

Experiments were carried out using *Arabidopsis thaliana* accession Columbia (Col‐0) as the wild‐type background. The *crtW* over‐expressors were obtained exploiting *Agrobacterium tumefaciens*‐mediated transformation of the 35S:PLS‐crtW construct in Col‐0, following the floral dip protocol (Clough and Bent, 1998). Transgenic seedlings were selected for resistance on kanamycin and subsequently verified by PCR for the presence of the transgene with the following primers: attB1 (GGGACAAGTTTGTACAAAAAAGCAGGCT) and PLS_crtW_Rv (TCAGGCGGTATCACCCTTAGT). Soil‐grown plants were cultivated at 23°C day/18°C night under neutral day photoperiod (12 light : 12 darkness) in single 6 cm pots using a 3:1 soil : perlite mixture (with HAWITA tray substrate), after seed vernalization at 4°C in the dark and germinated. The quantum irradiance was 80‐100 μmol photons m^‐2^ s^‐1^. For the *in vitro* selection of transformed plants, seeds were surface sterilized using 70% (v/v) ethanol and 10% (v/v) commercial bleach solution and then rinsed 5‐7 times with sterile distilled water. Seeds were then sown on solid sterile ½ strength MS medium [0.215% (w/v) Murashige‐Skoog (MS) salts (Sigma‐Aldrich), 0.8% agar (w/v), 0.5% (w/v) sucrose, pH 5.7]. Genomic DNA was extracted following the protocol described by Edwards et al., 1991.

### RNA extraction and gene expression analysis by qRT‐PCR

For qRT‐PCR analysis, total RNA was extracted from 4 weeks old plants as described by Kosmacz et al., 2015. cDNA was synthesized from 1 μg of total RNA using the Maxima Reverse Transcriptase kit (Life Technologies). Real‐time PCR amplification was performed on 12.5 ng cDNA with the ABI Prism 7300 sequence detection system (Applied Biosystems), using the PowerUp™ SYBR^®^ Green Master Mix (Applied Biosystems). Ubiquitin10 (*At4g053290*) was exploited as the housekeeping gene. A pair of specific primers was designed on the *crtW* gene sequence: sgCRTWfw (GGCACAACGCTCGTTCCTCT) and sgCRTWrv (AAACGCCACCAAGGCACAGT).

### Plasmid DNA Purification

Highly concentrated plasmid DNA required for protoplast transformation was extracted from 100‐mL bacterial culture (Luria‐Bertani medium) supplemented with the appropriate antibiotic. Bacterial pellets were extracted with an alkaline lysis protocol, according to Sambrook and Russell, 2001. Briefly, pellets were sequentially resupended in 2 mL of a buffer containing 50 mM Glucose, 25 mM Tris‐HCl and 10 mM EDTA (pH 8), lysed in 4 mL of a buffer containing 0.2 M NaOH and and 1% (w/v) SDS and neutralized in 3 mL of a buffer composed of 3 M potassium acetate in 11.5% (v/v) glacial acetic acid. Next, nucleic acids were precipitated with an equal volume of isopropanol and then treated with RNase A (Sigma‐Aldrich) for 1 h at 37°C. Following polyethylene glycol (PEG) precipitation (13% [w/v] PEG 8000 dissolved in 1.6 M NaCl), phenol : chloroform extraction was performed and nucleic acids were precipitated in absolute ethanol, with the aid of ammonium acetate 0.6M. Final elution was in nuclease‐free water.

### Protoplast isolation and transformation

Arabidopsis mesophyll protoplasts were isolated and transformed according to Wu et al., 2009. They were transfected with 4 µg of each effector plasmid and incubated in multiwell plates at 23°C in the dark for 16 hours.

### Light treatment of transfected protoplasts

Multiwell plates containing the transfected protoplasts were placed under led lamps, with the possibility to change both light wavelength and intensity. Red (615‐630 nm), green (515‐530 nm), blue (465‐480 nm) and white light were used in our experiments, testing an intensity range from 5 to 350 μmol m^−2^ s^−1^. For the reversion experiment the samples were transferred in darkness for 5 or 30 min before sampling.

### Luciferase Activity Quantification

Protoplasts were flash frozen in liquid nitrogen and lysed by adding 30 μl of Passive Lysis Buffer (Promega). NanoLuc activity was then measured using the Nano‐Glo^®^ Luciferase Assay (Promega) following the manufacturer’s instructions. Luminescence was detected with a tube Luminometer (Berthold). NanoLuc measurements were normalized according to the total protein concentration of each sample, determined by means of the Bio‐Rad colorimetric assay^®^, based on the Bradford dye‐binding method (Bradford, 1976).

### Extraction and quantification of carotenoid content

Pigments were extracted from leaves with 80% acetone buffered with Na2CO3 and quantified by reverse‐phase HPLC as described in Perozeni et al. 2020. In particular a Jasco LC‐4000 extrema HPLC system equipped with a C18 column (Synergi 4u Hydro‐RP 80A, Phenomenex, USA) was used. 15‐min gradient of ethyl acetate (0 to 100%) in acetonitrile‐water‐triethylamine (9:1:0.01, vol/vol/vol) at a flow rate of 1.5 ml/min was used (Lagarde, Beuf and Vermaas, 2000) and pigment detection was conducted with a Jasco 350–750 nm diode array detector. Keto‐carotenoids peaks were identified by comparing retention times and spectra to commercially available standards (CaroteNature GmbHas) as reported in Perozeni et al. 2020.

### Phenotypic Analysis

To quantify the phenotypic parameters of interest in *crtW* plants, pots were imaged at 17, 21 and 24 days after germination with a LabScanalyzer (LemnaTec, GmbH, Aachen, Germany), and images were taken and analysed as described by Ventura et al., 2020.

### Statistical analyses

Significant variations between genotypes or treatments were statistically evaluated using Student’s t‐test and ordinary one‐way ANOVA with multiple comparisons. Curves were fit to the data with linear/non‐linear regression. All the analysis were performed with GraphPad Prism 9 for Windows 10.

## Parsed Citations

Alcorta, J. et al. (2019) ‘Fischerella thermalis: a model organism to study thermophilic diazotrophy, photosynthesis and multicellularity in cyanobacteria’, Extremophiles. doi: 10.1007/s00792-019-01125-4. Google Scholar: Author Only Title Only Author and Title

Andres, J., Blomeier, T. and Zurbriggen, M. D. (2019) ‘Synthetic switches and regulatory circuits in plants’, Plant Physiology. doi: 10.1104/pp.18.01362. Google Scholar: Author Only Title Only Author and Title

Bai, C. et al. (2017) ‘Reconstruction of the astaxanthin biosynthesis pathwayin rice endosperm reveals a metabolic bottleneck at the level of endogenous β-carotene hydroxylase activity’, Transgenic Research, 26(1), pp. 13–23. doi: 10.1007/s11248-016-9977-x. Google Scholar: Author Only Title Only Author and Title

Bao, H. et al. (2017) ‘Additional families of orange carotenoid proteins in the photoprotective system of cyanobacteria’, Nature Plants, 3(July), pp. 1–11. doi: 10.1038/nplants.2017.89. Google Scholar: Author Only Title Only Author and Title

Bondanza, M. et al. (2020) ‘The Molecular Mechanisms of Photoactivation of Orange Carotenoid Protein Revealed by Molecular Dynamics’, pp. 1–8. doi: 10.1021/jacs.0c10461. Google Scholar: Author Only Title Only Author and Title

Bradford, M. M. (1976) ‘A rapid and sensitive method for the quantitation of microgram quantities of protein utilizing the principle of protein-dye binding’, Analytical Biochemistry. doi: 10.1016/0003-2697(76)90527-3. Google Scholar: Author Only Title Only Author and Title

Chatelle, C. et al. (2018) ‘A Green-Light-Responsive System for the Control of Transgene Expression in Mammalian and Plant Cells’, ACS Synthetic Biology, 7(5), pp. 1349–1358. doi: 10.1021/acssynbio.7b00450. Google Scholar: Author Only Title Only Author and Title

Choi, S. K. et al. (2005) ‘Characterization of β-carotene ketolases, CrtW, from marine bacteria by complementation analysis in Escherichia coli’, Marine Biotechnology, 7(5), pp. 515–522. doi: 10.1007/s10126-004-5100-z. Google Scholar: Author Only Title Only Author and Title

Clough, S. J. and Bent, A. F. (1998) ‘Floral dip: A simplified method for Agrobacterium-mediated transformation of Arabidopsis thaliana’, Plant Journal. doi: 10.1046/j.1365-313X.1998.00343.x. Google Scholar: Author Only Title Only Author and Title

Corpet, F. (1988) ‘Multiple sequence alignment with hierarchical clustering’, Nucleic Acids Research. doi: 10.1093/nar/16.22.10881. Google Scholar: Author Only Title Only Author and Title

DeLano, W. L. (2020) ‘The PyMOL Molecular Graphics System, Version 2.3’, Schrödinger LLC.

Dixon, A. S. et al. (2016) ‘NanoLuc Complementation Reporter Optimized for Accurate Measurement of Protein Interactions in Cells’, ACS Chemical Biology, 11(2), pp. 400–408. doi: 10.1021/acschembio.5b00753. Google Scholar: Author Only Title Only Author and Title

Dominguez-Martin, M. A. and Kerfeld, C. A. (2019) ‘Engineering the orange carotenoid protein for applications in synthetic biology’ Current Opinion in Structural Biology, 57, pp. 110–117. doi: 10.1016/j.sbi.2019.01.023. Google Scholar: Author Only Title Only Author and Title

Edwards, K., Johnstone, C. and Thompson, C. (1991) ‘A simple and rapid method for the preparation of plant genomic DNAfor PCR analysis’, Nucleic Acids Research. doi: 10.1093/nar/19.6.1349. Google Scholar: Author Only Title Only Author and Title

Gupta, S. et al. (2015) ‘Local and global structural drivers for the photoactivation of the orange carotenoid protein’, Proceedings of the National Academy of Sciences of the United States of America, 112(41), pp. E5567–E5574. doi: 10.1073/pnas.1512240112. Google Scholar: Author Only Title Only Author and Title

Gutmann, A. (2011) ‘The ethics of synthetic biology: Guiding principles for emerging technologies’, Hastings Center Report. doi: 10.1002/j.1552-146X.2011.tb00118.x. Google Scholar: Author Only Title Only Author and Title

Hall, M. P. et al. (2012) ‘Engineered luciferase reporter from a deep sea shrimp utilizing a novel imidazopyrazinone substrate’, ACS Chemical Biology, 7(11), pp. 1848–1857. doi: 10.1021/cb3002478. Google Scholar: Author Only Title Only Author and Title

Hart, J. E. et al. (2019) ‘Engineering the phototropin photocycle improves photoreceptor performance and plant biomass production’, Proceedings of the National Academy of Sciences of the United States of America, 116(25), pp. 12550–12557. doi: 10.1073/pnas.1902915116. Google Scholar: Author Only Title Only Author and Title

Karimi, M., Inze, D. and Depicker, A. (2002) ‘GATEWAYTM vectors for Agrobacterium-mediated plant transformation’, Trends in Plant Science.

Kay Holt, T. and Krogmann, D. W. (1981) ‘A carotenoid-protein from cyanobacteria’, BBA-Bioenergetics. doi: 10.1016/0005-2728(81)90045-1. Google Scholar: Author Only Title Only Author and Title

Kennedy, M. J. et al. (2010) ‘Rapid blue-light-mediated induction of protein interactions in living cells’, Nature Methods, 7(12), pp. 973–975. doi: 10.1038/nmeth.1524. Google Scholar: Author Only Title Only Author and Title

Kerfeld, C. A. et al. (2003) ‘The Crystal Structure of a Cyanobacterial Water-Soluble Carotenoid Binding Protein let chlorophyll, deexciting it in a preemptive reaction before singlet oxygen can be formed (reviewed in [1–3]). For example, transforming cyanobacteria with various’, Structure, 11(02), pp. 55–65. Google Scholar: Author Only Title Only Author and Title

Kirilovsky, D. and Kerfeld, C. A. (2013) ‘The Orange Carotenoid Protein: A blue-green light photoactive protein’, Photochemical and Photobiological Sciences, 12(7), pp. 1135–1143. doi: 10.1039/c3pp25406b. Google Scholar: Author Only Title Only Author and Title

Kosmacz, M. et al. (2015) ‘The stability and nuclear localization of the transcription factor RAP2.12 are dynamically regulated byoxygen concentration’, Plant, Cell and Environment. doi: 10.1111/pce.12493. Google Scholar: Author Only Title Only Author and Title

Lagarde, D., Beuf, L. and Vermaas, W. (2000) ‘Increased production of zeaxanthin and other pigments by application of genetic engineering techniques to Synechocystis sp. strain PCC 6803’, Applied and Environmental Microbiology. doi: 10.1128/AEM.66.1.64-72.2000. Google Scholar: Author Only Title Only Author and Title

Lechno-Yossef, S. et al. (2017) ‘Synthetic OCP heterodimers are photoactive and recapitulate the fusion of two primitive carotenoproteins in the evolution of cyanobacterial photoprotection’, Plant Journal, 91(4), pp. 646–656. doi: 10.1111/tpj.13593. Google Scholar: Author Only Title Only Author and Title

Leverenz, R. L. et al. (2014) ‘Structural and functionalmodularity of the orange carotenoid protein: Distinct roles for the N- and C-terminal domains in cyanobacterial photoprotection’, Plant Cell, 26(1), pp. 426–437. doi: 10.1105/tpc.113.118588. Google Scholar: Author Only Title Only Author and Title

Leverenz, R. L. et al. (2015) ‘A12 Å carotenoid translocation in a photoswitch associated with cyanobacterial photoprotection’, Science, 348(6242), pp. 1463–1466. doi: 10.1126/science.aaa7234. Google Scholar: Author Only Title Only Author and Title

Merezhko, M. et al. (2020) ‘Live-cell monitoring of protein localization to membrane rafts using protein-fragment complementation’, Bioscience Reports, 40(1), pp. 1–13. doi: 10.1042/BSR20191290. Google Scholar: Author Only Title Only Author and Title

‘Method of the Year 2010’ (2011) Nature Methods. doi: 10.1038/nmeth.f.321. Google Scholar: Author Only Title Only Author and Title

Misawa, N. et al. (1995) ‘Canthaxanthin biosynthesis by the conversion of methylene to keto groups in a hydrocarbon β-carotene by a single gene’, Biochemical and Biophysical Research Communications. doi: 10.1006/bbrc.1995.1579. Google Scholar: Author Only Title Only Author and Title

Möglich, A. et al. (2010) ‘Structure and function of plant photoreceptors’, Annual Review of Plant Biology. doi: 10.1146/annurev-arplant-042809-112259. Google Scholar: Author Only Title Only Author and Title

Müller, K. et al. (2013) ‘Multi-chromatic control of mammalian gene expression and signaling’, Nucleic Acids Research, 41(12). doi: 10.1093/nar/gkt340. Google Scholar: Author Only Title Only Author and Title

Nielsen, A. Z. et al. (2013) ‘Redirecting photosynthetic reducing power toward bioactive natural product synthesis’, ACS Synthetic Biology, 2(6), pp. 308–315. doi: 10.1021/sb300128r. Google Scholar: Author Only Title Only Author and Title

Nisar, N. et al. (2015) ‘Carotenoid metabolism in plants’, Molecular Plant, 8(1), pp. 68–82. doi: 10.1016/j.molp.2014.12.007. Google Scholar: Author Only Title Only Author and Title

Olson, E. J. and Tabor, J. J. (2014) ‘Optogenetic characterization methods overcome key challenges in synthetic and systems biology’, Nature Chemical Biology. doi: 10.1038/nchembio.1559. Google Scholar: Author Only Title Only Author and Title

Paik, I. and Huq, E. (2019) ‘Plant photoreceptors: Multi-functional sensory proteins and their signaling networks’, Seminars in Cell and Developmental Biology. doi: 10.1016/j.semcdb.2019.03.007. Google Scholar: Author Only Title Only Author and Title

Perozeni, F. et al. (2020) ‘Turning a green alga red: engineering astaxanthin biosynthesis by in tragenic pseudogene revival in Chlamydomonas reinhardtii’, Plant Biotechnology Journal. doi: 10.1111/pbi.13364. Google Scholar: Author Only Title Only Author and Title

Pouvreau, B. et al. (2020) ‘A Versatile High Throughput Screening Platform for Plant Metabolic Engineering Highlights the Major Role of ABI3 in Lipid Metabolism Regulation’, Frontiers in Plant Science. doi: 10.3389/fpls.2020.00288. Google Scholar: Author Only Title Only Author and Title

Punginelli, C. et al. (2009) ‘Influence of zeaxanthin and echinenone binding on the activity of the Orange Carotenoid Protein’, Biochimica et Biophysica Acta - Bioenergetics, 1787(4), pp. 280–288. doi: 10.1016/j.bbabio.2009.01.011. Google Scholar: Author Only Title Only Author and Title

Ruiz-Sola, M. Á. and Rodríguez-Concepción, M. (2012) ‘Carotenoid Biosynthesis in Arabidopsis: A Colorful Pathway’ The Arabidopsis Book, 10, p. e0158. doi: 10.1199/tab.0158. Google Scholar: Author Only Title Only Author and Title

Sambrook, J. and Russell, D. W. (2001) ‘Molecular Cloning: ALaboratory Manual, Third Edition’, in Molecular Cloning: a laboratory a manual,.

Sineshchekov, O. A., Jung, K. H. and Spudich, J. L. (2002) ‘Two rhodopsins mediate phototaxis to low- and high-intensity light in Chlamydomonas reinhardtii’, Proceedings of the National Academy of Sciences of the United States of America. doi: 10.1073/pnas.122243399. Google Scholar: Author Only Title Only Author and Title

Sun, T. et al. (2018) ‘Carotenoid Metabolism in Plants: The Role of Plastids’, Molecular Plant. doi: 10.1016/j.molp.2017.09.010. Google Scholar: Author Only Title Only Author and Title

Tilbrook, K. et al. (2013) ‘The UVR8 UV-B Photoreceptor: Perception, Signaling and Response’, The Arabidopsis Book. doi: 10.1199/tab.0164. Google Scholar: Author Only Title Only Author and Title

Ventura, I. et al. (2020) ‘Arabidopsis phenotyping reveals the importance of alcohol dehydrogenase and pyruvate decarboxylase for aerobic plant growth’, Scientific Reports. doi: 10.1038/s41598-020-73704-x. Google Scholar: Author Only Title Only Author and Title

Verhounig, A., Karcher, D. and Bock, R. (2010) ‘Inducible gene expression from the plastid genome by a synthetic riboswitch’, Proceedings of the National Academy of Sciences of the United States of America, 107(14), pp. 6204–6209. doi: 10.1073/pnas.0914423107. Google Scholar: Author Only Title Only Author and Title

Wang, F. Z. et al. (2020) ‘Split Nano luciferase complementation for probing protein-protein interactions in plant cells’, Journal of Integrative Plant Biology, 62(8), pp. 1065–1079. doi: 10.1111/jipb.12891. Google Scholar: Author Only Title Only Author and Title

Webb, B. and Sali, A. (2016) ‘Comparative protein structure modeling using MODELLER’, Current Protocols in Bioinformatics. doi: 10.1002/cpbi.3. Google Scholar: Author Only Title Only Author and Title

Wilson, A. et al. (2006) ‘A soluble carotenoid protein involved in phycobilisome-related energy dissipation in cyanobacteria’, Plant Cell, 18(4), pp. 992–1007. doi: 10.1105/tpc.105.040121. Google Scholar: Author Only Title Only Author and Title

Wilson, A. et al. (2008) ‘A photoactive carotenoid protein acting as light intensity sensor’, Proceedings of the National Academy of Sciences of the United States of America, 105(33), pp. 12075–12080. doi: 10.1073/pnas.0804636105. Google Scholar: Author Only Title Only Author and Title

Wilson, A. et al. (2011) ‘Essential role of two tyrosines and two tryptophans on the photoprotection activity of the Orange Carotenoid Protein’, Biochimica et Biophysica Acta - Bioenergetics, 1807(3), pp. 293–301. doi: 10.1016/j.bbabio.2010.12.009. Google Scholar: Author Only Title Only Author and Title

Wu, F. H. et al. (2009) ‘Tape-arabidopsis sandwich - A simpler arabidopsis protoplast isolation method’, Plant Methods. doi: 10.1186/1746-4811-5-16. Google Scholar: Author Only Title Only Author and Title

Wu, Y. P. and Krogmann, D. W. (1997) ‘The orange carotenoid protein of Synechocystis PCC 6803’, Biochimica et Biophysica Acta - Bioenergetics, 1322(1), pp. 1–7. doi: 10.1016/S0005-2728(97)00067-4. Google Scholar: Author Only Title Only Author and Title

Xiang, N. et al. (2020) ‘Erratum: Using synthetic biology to overcome barriers to stable expression of nitrogenase in eukaryotic organelles (Proceedings of the National Academy of Sciences of the United States of America (2020) 117 (16537-16545) DOI: 10.1073/pnas.2002307117)’, Proceedings of the National Academy of Sciences of the United States of America, 117(39), p. 24602. doi: 10.1073/pnas.2017731117. Google Scholar: Author Only Title Only Author and Title

Yazawa, M. et al. (2009) ‘Induction of protein-protein interactions in live cells using light’, Nature Biotechnology, 27(10), pp. 941–945. doi: 10.1038/nbt.1569. Google Scholar: Author Only Title Only Author and Title

Yoo, S. D., Cho, Y. H. and Sheen, J. (2007) ‘Arabidopsis mesophyll protoplasts: A versatile cell system for transient gene expression analysis’, Nature Protocols. doi: 10.1038/nprot.2007.199. Google Scholar: Author Only Title Only Author and Title

Zhong, Y. J. et al. (2011) ‘Functional characterization of various algal carotenoid ketolases reveals that ketolating zeaxanthin efficiently is essential for high production of astaxanthin in transgenic Arabidopsis’, Journal of Experimental Botany. doi: 10.1093/jxb/err070. Google Scholar: Author Only Title Only Author and Title

